# Identification of a new orthonairovirus associated with human febrile illness in China

**DOI:** 10.1101/2020.10.24.353458

**Authors:** Jun Ma, Xiao-Long Lv, Xu Zhang, Shu-Zheng Han, Ze-Dong Wang, Liang Li, He-Ting Sun, Li-Xin Ma, Zheng-Lei Cheng, Jian-Wei Shao, Chen Chen, Ying-Hua Zhao, Liyan Sui, Lin-Na Liu, Jun Qian, Wei Wang, Quan Liu

## Abstract

The genus *Orthonairovirus* of the family *Nairoviridae* includes the important tick-transmitted pathogens, Crimean-Congo hemorrhagic fever virus (CCHFV) and Nairobi sheep disease virus (NSDV), as well as many other poorly characterized viruses isolated from ticks, birds, and mammals^1,2^. Here we identified a novel orthonairovirus, designated Sōnglǐng virus (SGLV), from patients who reported being bitten by a tick in China. The genome of SGLV shared similar structural features with orthonairoviruses, with 46.5–65.7% sequence identify. Phylogenetic analysis showed that SGLV belonged to the *Tamdy orthonairovirus* and formed a unique clade in the *Nairoviridae* family. Electron microscopy revealed typical morphological characteristics of orthonairoviruses. The isolated SGLVs from the blood samples of patients could induce cytopathic effects in human hepatoma cells. SGLV infection was confirmed in 42 patients in 2017-2018, with the main clinical manifestations of headache, fever, depression, fatigue and dizziness. Serological assays showed that 69% patients generated virus-specific antibody responses in the acute phase. In contrast, neither SGLV viral RNA nor specific antibodies against SGLV were detected in healthy individuals. SGLV was also detected in *Ixodes crenulatus, Haemaphysalis longicornis, Haemaphysalis concinna*, and *Ixodes persulcatus* in northeastern China. Collectively, a newly discovered orthonairovirus was shown to be associated with human febrile illness in northeastern China.

---

Tick-borne orthonairoviruses have been considered as a global health threat to humans and animals^3^. CCHFV causes fatal hemorrhagic fever in humans in Asia, Africa, and Europe^4,5^, and NSDV leads to hemorrhagic gastroenteritis in sheep and goats in Africa and Asia^6^. Other orthonairoviruses, such as Dugbe virus, Tǎchéng tick virus 1, Erve virus, Tamdy virus, and Soldado virus, can cause fever, headache, rashes, pruritus, thrombocytopenia, or neurological disorders in humans^7–11^. Here we identified a newly discovered orthonairovirus, designated Sōnglǐng virus (SGLV), associated with human febrile illness in China.

In April 2017, a 47-year-old male farmer from the Sōnglǐng town 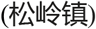 in Hēilóngjiāng Province 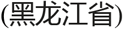 presented to a hospital in Hulunbuir/Hūlúnbèi’ěr 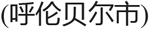 of Inner Mongolia, China, with fever, headache, fatigue, and depression. The tick-borne pathogens, such as tick-borne encephalitis virus (TBEV), severe fever with thrombocytopenia syndrome virus (SFTSV), Alongshan virus (ALSV), *Borrelia burgdorferi* sensu lato, *Borrelia miyamotoi, Anaplasma* spp., *Babesia* spp., and *Rickettsia* spp., were tested negative^12–15^. Blood specimens were used for viral metagenomic analysis, producing 1,522 virus-related reads (Fig. 1a), 68% of which were assembled into 29 contigs targeting Wēnzhōu tick virus (Supplementary Table 1–3).

**Fig. 1.**
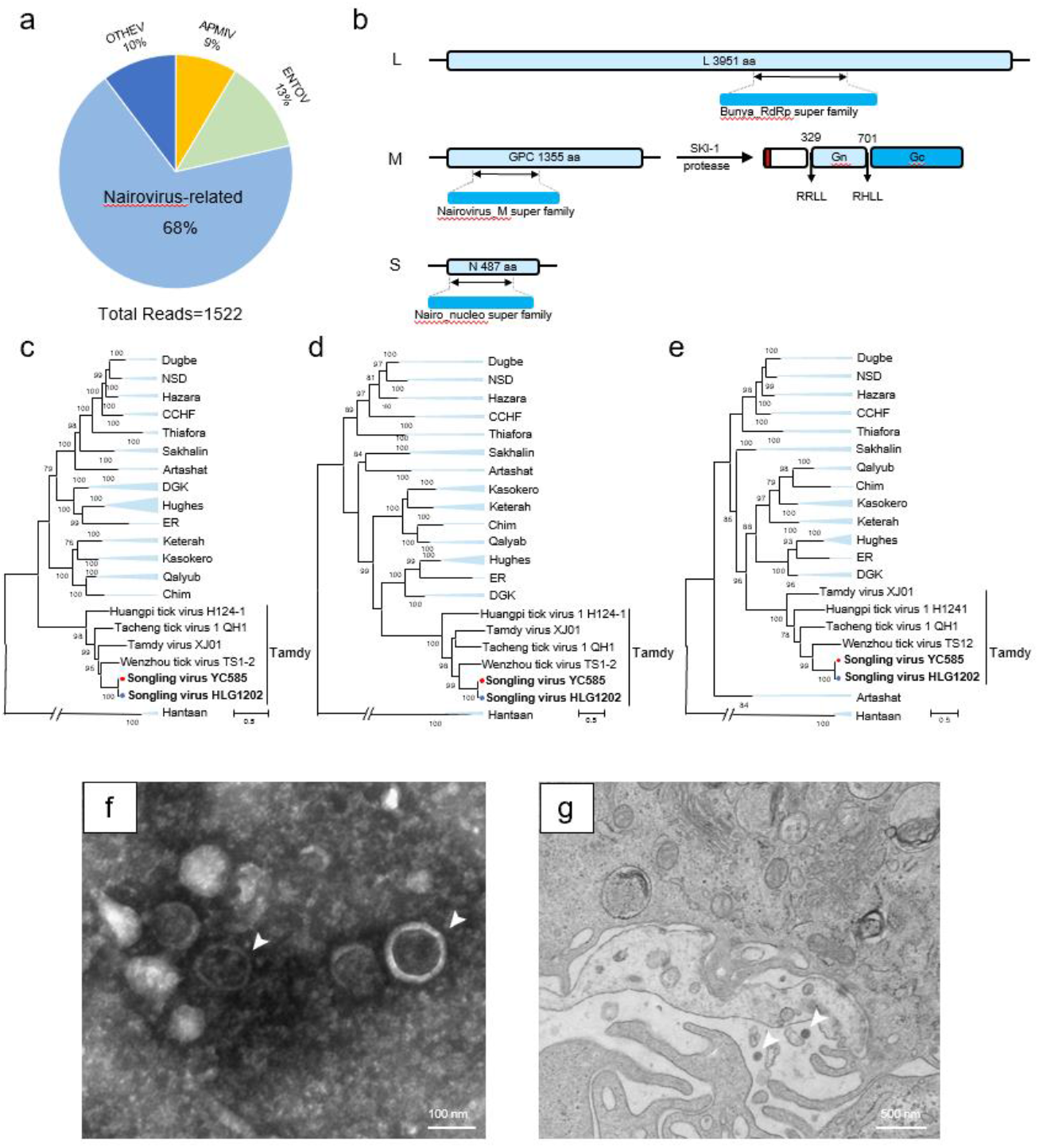
Discovery and characterization of SGLV. **a,** A pie chart illustrating the composition of virus-related reads from metagenomic analysis of the index patient’s blood sample. Percentages of the virus-related reads for orthonairovirus, Enterovirus B (ENTOV), Acanthamoeba polyphaga mimivirus (APMIV), and other viruses (OTHEV) are shown. **b,** Genome organization and putative coding regions of SGLV. The viral genome includes large (L), medium (M), and small (S) segments that contained the predicted RNA-directed RNA polymerase (RdRp), glycoprotein precursor (GPC), and nucleoprotein (N), respectively. The super families of their encoding proteins are shown in blue boxes. In the M segment, the subtilisin/kexin-isozyme-1 (SKI-1) protease cleavage sites and sequences of GPC are indicated with black arrows, and the signal peptide is marked with red. **c–e,** Phylogenetic analysis of SGLVs. The trees were constructed based on the complete amino acid sequences of RdRp (**c**), GPC (**d**), and N (**e**) with maximum likelihood method. Viruses representing *Artashat, Chim, Crimean-Congo hemorrhagic fever (CCHF), Dera Ghazi Khan (DGK), Dugbe, Estero Real (ER), Hazara, Hughes, Kasokero, Keterah, Nairobi sheep disease* (NSD), *Qalyub, Sakhalin, Tamdy*, and *Thiafora Orthonairoviruses* were included in the analysis. SGLVs isolated in this study are marked with a blue dot. Detailed information of involved orthonairoviruses are shown in Supplementary Table 4. **f,** Negatively stained virions (arrows) purified from infected SMMC-7721 cells. **g,** Transmission electron microscopy showed virions (arrows) in the cytoplasm of infected SMMC-7721 cells.

The complete genome of the virus (strain HLJ1202), determined by overlap RT-PCR and rapid amplification of cDNA ends (Extended data Fig. 1), included large (L, 12,001 nt), medium (M, 4,335 nt), and small (S, 1,881 nt) segments (GenBank accession numbers MT328776, MT328775, and MT328777). This novel virus was tentatively named Sōnglǐng virus (SGLV), which had similar genomic structure with typical orthonairoviruses, such as CCHFV, Hughes virus, Erve virus, and Tamdy virus, with three segments encoding the large (L) protein, glycoprotein precursor (GPC), and nucleoprotein (N), respectively (Fig. 1b, Extended data Fig. 2a)^2^. The 5’-terminus of SGLV L and S segments had a typical terminal sequence UCUCAAAGA of orthonairoviruses, while the 3’-terminus of all three segments ended up with CCCATGT^2^, and the terminus of M segment had a reverse-complementary sequence ACATGGG (Extended data Fig. 2b).

The L segment encoded a large protein of 3,951 amino acids (aa). The predicted catalytic polymerase domain was located at aa 2279-2594, with the six conserved motifs (pre-motif A and motifs A-E)^2^. Others motifs, including ovarian tumor-like domain, zinc finger, and leucine zipper motifs, were also presented in SGLV (Extended data Fig. 3)^16,17^. The M segment encoded a 1,335 aa glycoprotein precursor (Gn and Gc). The conserved N-terminal signal peptide followed by a highly O-glycosylated mucin-like domain was discovered in GPC of SGLV.^18^ Moreover, the cleavage sites of the subtilisin/kexin-isozyme-1 (SKI-1) protease, which cleaved GPC into mature Gn and Gc, were identified at the sites RRLL_329_↓ and RHLL_701_↓, respectively (Fig. 1b)^19^. The S segment encoded a 487 aa nucleoprotein, similar to those of other orthonairoviruses (Extended data Fig. 2a)^20^. A positively charged region involved in RNA binding was identified at the C-terminus of SGLV N protein (Extended data Fig. 4)^21,22^.

We also obtained partial sequences of three other viral strains (HLB178, HLJ1175, and GH1185), which were genetically related to the strain HLJ1202. Phylogenetic analysis showed that SGLVs were grouped into the Tamdy genogroup and formed a unique clade distinct from other *Tamdy orthonairoviruses*, indicating that SGLV may represent a new species in the family of *Nairoviridae* (Fig. 1c-e, Extended data Fig. 5). Comparison of the nucleotides and amino acids demonstrated that SGLV was more closely related to the Tamdy orthonairoviruses (46.5–65.7% sequence similarity) than to other genogroups of orthonairoviruses, with 24.6–50.6% sequence identity (Extended data Fig. 6, Supplementary Table 5–6), further confirming that SGLV is a new species in the *Tamdy Orthonairovirus*. With the exception of Burana virus, all these Tamdy orthonairoviruses have been detected in ticks in China, suggesting that Tamdy orthonairoviruses have a high genetic diversity in China^8,23,24^.

From April 7, 2017 to December 6, 2018, a total of 42 from 658 hospitalized patients who had a history of tick bite were confirmed to be infected with SGLV by real-time RT-PCR (Supplementary Table 7, Extended data Fig. 7). Of them, 12 were found in 2017 (12/348) and 30 were in 2018 (30/310); 29 were from Inner Mongolia and 13 were from Heilongjiang (Extended data Fig. 8). The infection usually occurred between May and July (90.5%). All the patients were field workers or lived in forestry areas, the median age of the 42 cases was 48 years old, ranging from 24 to 70 years old; 26 (61.9%) were 40–60 years of age, 29 (69.0%) were male. The incubation period of SGLV infection ranged from 1 to 25 days (median, 6 days; interquartile range, 4–8 days).

Headache (76%) and fever (74%) were the most common clinical manifestations, and 52% patients reported depression and 50% reported fatigue or dizziness. Other clinical findings included myalgia or arthralgia, rash or petechiae, anorexia, and nausea. A few cases showed vomiting, chest tightness, chilling, cough, tinnitus, stomach discomfort, diarrhea, malaise, insomnia, coma, abdominal pain, or tenderness (Table 1). These clinical manifestations have also been found in other tick-borne diseases, such as tick-borne encephalitis virus, Alongshan virus, anaplasmosis, babesiosis, and rickettsiosis^12–15^. Thus, SGLV should be included in the differential diagnosis from other tick-borne pathogens.

**Table 1.** Clinical manifestations of SGLV-infected patients

| Clinical manifestation | Confirmed cases (n=42) |
| --- | --- |
| Headache | 32 (76%) |
| Fever | 31 (74%) |
| Depression | 22 (52%) |
| Fatigue | 21 (50%) |
| Dizziness | 21 (50%) |
| Myalgia or arthralgia | 14 (33%) |
| Rash or petechiae | 13 (31%) |
| Anorexia | 13 (31%) |
| Nausea | 11 (26%) |
| Vomiting | 7 (17%) |
| Chest tightness | 6 (14%) |
| Chilling | 4 (10%) |
| Cough | 4 (10%) |
| Tinnitus | 3 (7%) |
| Stomach discomfort | 3 (7%) |
| Diarrhea | 2 (5%) |
| Malaise | 2 (5%) |
| Insomnia | 2 (5%) |
| Coma | 1 (2%) |
| Abdominal pain or tenderness | 1 (2%) |

Laboratory abnormalities included elevated levels of homocysteine (93%) and high-sensitivity C-reactive protein (82%), decreased level of hypersensitive troponin (83%), myoglobin (83%), and apolipoprotein (78%), which may be related to inflammation, coagulation, or endothelial dysfunction^25,26^. Liver injury was indicated by the enhanced serum aspartate aminotransferase (32%) and alanine aminotransferase (16%) (Supplementary Table 8).

All these patients were empirically treated with a combination of benzylpenicillin sodium (4 million units per day) and ribavirin (0.5 g per day). The duration of the symptoms ranged from 2 days to 60 days (median, 12 days; interquartile range, 9–17 days). The symptoms usually resolved in 2–27 days after treatment (Fig. 2a).

**Fig. 2.**
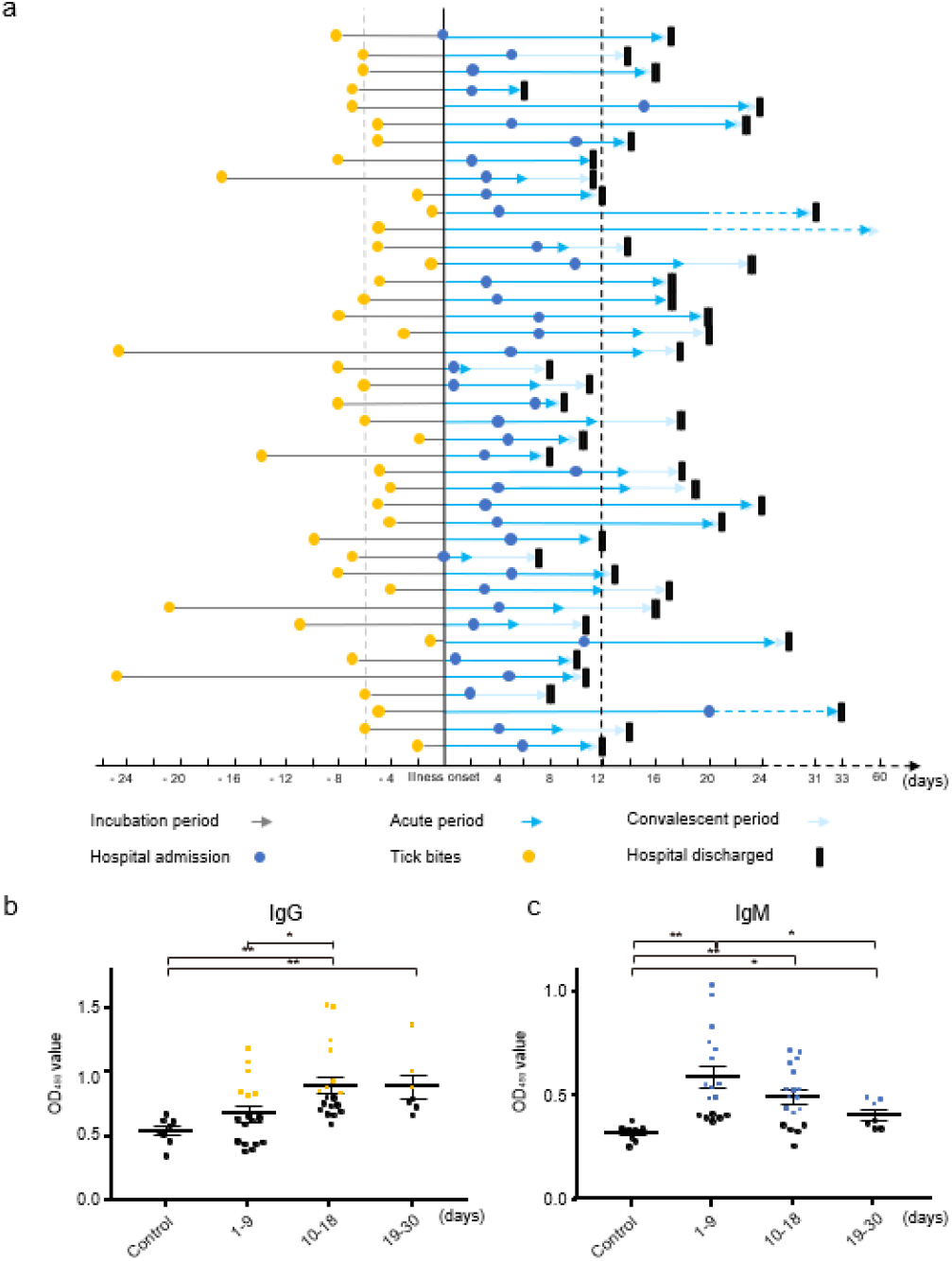
A timeline of patients with SGLV infection and serological investigation. **a,** Timeline of patients with SGLV infection. Patients were ordered in chronological order based on the date of sample collection. We defined headache and fever as the cutoff of the acute SGLV disease. Different periods of disease course are indicated with different-colored arrows and various milestones are presented with different-colored symbols. The median of incubation and acute periods are presented with gray dot and black dot lines, respectively. **b,** Presentation of OD450 values for ELISA IgM of serum samples from the SGLV-infected patients. The cut-off value was 0.5 and positive samples were marked with blue. **c,** Presentation of OD450 values for ELISA IgG of serum samples from the SGLV-infected patients. The cut-off value was 0.8 and positive samples were marked with yellow. Control group consisted of 8 pools of healthy individuals, with 12-13 healthy serum samples in each pool. Mean and standard error of mean (SEM) are indicated for each group and differences between each group are considered statistically significant at p-values <0.05 and marked with asterisks (\**p* < 0.05, \*\**p* < 0.01).

SGLV was isolated from the blood samples of index patient HLJ1202 and three other patients (HLB178, HLJ1175, and GH1185). Virus-induced cytopathogenic effects (CPE) were observed in the infected SMMC-7721 cells, but not in Vero and BHK-21 cells. SGLV could be detected in the three cell lines using IFA (Extended data Fig. 10). Electron micrographs showed that negative-stained viral particles were generally enveloped spheres, with some surface projections and a diameter ranging from approximately 80 to 100 nm (Fig. 1f). Viral particles were observed in the ultrathin sections of SGLV-infected cells (Fig. 1g). This virus morphology consisted with the *Nairoviridae* family^27^. The virus was isolated from the blood samples of SGLV-infected patients, thus, may present a potential blood transfusion safety risk^28^.

We evaluated serological reaction against SGLV in infected patients by using indirect ELISA based on recombinant nucleoprotein (Extended data Fig. 9) and microneutralization. The ELISA results showed an IgG positive rate of 61.9% (26/42) and an IgM positive rate of 40.4% (17/42). Neutralizing antibodies were detected a prevalence of 26.1% (11/42). In total, 29 (69%) of 42 patients generated virus-specific antibody responses in the acute phase (Supplementary Table 9), but the control had no serological reaction. We further classified all serum samples into three groups according to the sampling time after tick bite (Fig. 2b), showing that the average IgM titer was higher in the 1–9 days group, with a downtrend appeared when compared with 10–18 days and 18–30 days groups. An uptrend of the average IgG titer was observed during the acute phase of SGLV infection. However, neither viral RNA nor specific antibodies against SGLV was detected in the healthy individuals.

As all patients had tick bites before the illness onset, we investigated the presence of SGLV in ticks collected in the case-distributed areas by real-time RT-PCR. Four tick species, including *Ixodes crenulatus, Ixodes persulcatus*, *Haemaphysalis concinna*, and *Haemaphysalis longicornis*, were tested positive for SGLV, with a natural infection rate of 0.8–5.5% (Supplementary Table 10). The highest infection rate of SGLV infection was observed in *I. crenulatus* (5.5%) that is widely distributed in Asia and Europe, suggesting that *I. crenulatus* may be the main carrier of the virus. However, their ability to transmit SGLV needs to be experimentally verified. These viruses from ticks were closely related to those in blood samples from the tick-bitten patients (Fig. 3). Thus, SGLV infection should be suspected in individuals who have been exposed to tick bites in these areas.

**Fig. 3.**
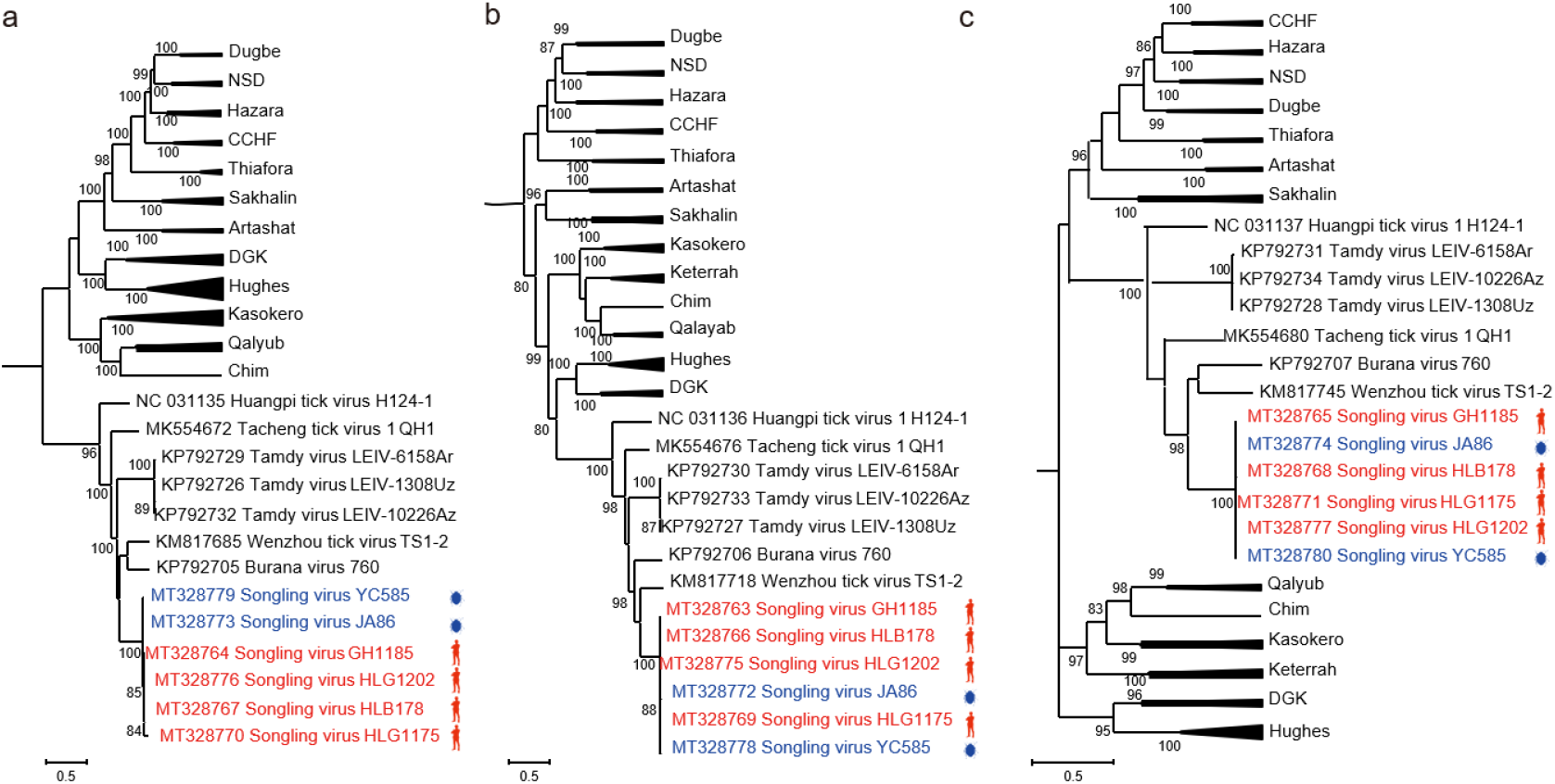
Phylogenetic analysis of SGLV from humans and ticks. The phylogenetic trees were generated based on the RNA-directed-RNA-polymerase (**a,** aa 2182-2446), glycoprotein precursor (**b,** aa 258-617) and nucleoprotein (**c,** aa 59-296) with maximum likelihood method. SGLVs isolated from humans and ticks are labeled with red and blue, respectively. The information of viruses used in the analysis are provided in Supplementary Tables 4 and 11. CCHF, Crimean-Congo hemorrhagic fever; NSD, Nairobi sheep disease; DGK, Dera Ghazi Khan; Chim, Chim virus.

Taken together, the newly discovered orthonairovirus was identified in patients with a tick bite history in northeastern China. Our findings provide evidence of an association between the febrile illness and this novel orthonairovirus. However, only a few patients with SGLV infection were identified in this study, thus, an active surveillance should be conducted to identify more patients and to determine the clinical characteristics of this emerging virus in infected patients. The present study was not designed to fulfill the Koch’s postulates of identifying a novel etiology. An animal infection model should further be developed to simulate the clinical symptoms of patients with SGLV infection, as the Koch’s postulates are considered a sufficient but not a necessary condition to establish causation in microbe studies^29,30^. The virus should be included in the differential diagnosis of tick bite patients, and the public health significance of the emerging virus necessitates further investigation in a larger geographical area.

## Methods

### Study design and sample collection

From April, 2017 to December, 2018, we recruited hospitalized patients with a history of tick bite to detect potential tick-borne pathogens at the Inner Mongolia General Forestry Hospital (120°44’36.82” E, 49°16’57.07” N), Yakeshi, Inner Mongolia Autonomous Region in northeastern China. The hospital is one of the largest in Hulunbuir city, which has a population of 2.5 million, lies between grasslands and forested highlands. A standardized questionnaire was administrated to each patient to obtain data about demography, medical history, and tick exposure. Clinical manifestations and laboratory tests were retrieved from medical records.

Confirmed cases were defined as patients who were tested positive for SGLV viral RNA in a blood sample by real-time RT-PCR. Patients who had co-infection with other known tick-borne pathogens were excluded (Supplementary Table 12). In addition, serum samples from 100 healthy individuals who lived in the same areas and never had tick bite were also collected during the same period.

Ticks were collected by flagging vegetation in Inner Mongolia (120°46’43.19”-123°48’28.41”E, 46°46’14.02”-51°15’15.62”N), Hēilóngjiāng (122°19’59.99”-130°41’23.11”E, 45° 5’23.58”-52°57’1.36”N), and Jílín (128°13’56.71”E, 43°22’22.94”N) provinces of China in May, 2015, and identified to species.

### Metagenomic analysis

Viral metagenomic analysis was conducted as previously described^31^. Briefly, total RNA extracted from the blood of index patient was digested by DNase I, and the rRNA was removed with Ribo-Zero Magnetic Gold Kit (Epicentre, USA). The extracted RNA was reverse transcribed with anchored random primers, and the double-stranded cDNA (dscDNA) was synthesized with Klenow fragment. The dscDNA was amplified by the sequence-independent single primer, followed by paired-end (150-bp) sequencing on novaseq 6000 platform (Illumina, USA) at the Beijing Genome Institute (BGI, China)^12^. Sequencing reads were demultiplexed and trimmed with Trimmomatic^32^. The resulting reads were blasted against the nonredundant protein (nr) database by using BLASTX, before de novo assembly using Megahit^33^. Virus-associated contigs were mapped against their corresponding reference sequences. The successfully assembled orthonairovirus-related contigs were further verified by RT-PCR. The primers are available upon request.

### Genome determination and phylogenetic analysis

The complete genomes of SGLV were obtained using overlap RT-PCR with the specific primers based on the above assembled contigs. The terminal sequences were recovered using a SMARTer^®^ RACE 5’/3’ Kit (Takara, Japan) according to the manufacturer’s protocol. The sequencing strategy of complete genome of SGLV is shown in Extended data Fig. 10, and the used primers are listed in Supplementary Table 13. The PCR products were cloned into the pM18-T vector (Takara) and sequenced.

Viral genomes obtained from this study were aligned with the known orthonairoviruses by using MAFFT version 7 with the E-INS-I algorithm^34,35^. After analyzing the suitable model by Prottest 3.4^36^, phylogenetic trees were generated using the maximum likelihood (ML) method in PhyML version 3 o 0 with LG model and a Subtree Pruning and Regrafting (SPR) topology searching algorithm^37^.

### Diagnostic tests

To detect the SGLV infection, total RNA from 140 μl patients’ serum or blood samples was extracted by using QIAamp Viral RNA Mini Kit (Qiagen, German) and then reverse-transcribed with PrimeScript™ RT reagent Kit with gDNA Eraser (TaKaRa). The specific primers targeting S segment of SGLV were designed for diagnostic real-time RT-PCR assay (Forward: 5’-ATGGCACCTGTGTATGAG-3’; Reverse: 5’-AGGCTTTCGTACTCCTTG-3’). The 20 μl reaction mixture contained 10 μl 2×TB green Premix DimerEraser (Takara), 1 μl of each primer, 6 μl of RNase-free H_2_O, and 2 μl of cDNA template. Amplification was conducted as follow: 95 °C for 30 s, 40 cycles of 95°C for 5 s, 60°C for 30 s, and 72°C for 30 s. Based on the lower limit of standard curve, the sample was considered positive if the cycle threshold (Ct) less than 36.8.

### Virus isolation

Human hepatocellular carcinoma (SMMC-7721), African green monkey kidney (Vero), and baby hamster kidney (BHK-21) cells were used for virus isolation. They were cultured in Dulbecco’s modified Eagle’s medium (DMEM; Gibco, USA) supplemented with 10% fetal bovine serum (Gibco, USA) and 1% penicillin-streptomycin in 5% CO_2_ at 37°C. The blood samples from infected patients were centrifuged at 12,000 *g* for 10 min, and the supernatant was diluted 10 times with DMEM before inoculated onto the confluent monolayer of SMMC-7721, Vero, and BHK-21 cells, respectively. After a 2-hour incubation at 37°C, the cells were washed with phosphate-buffered saline and incubated at 37°C, followed by monitoring daily for cytopathic effects (CPE). After three rounds of blinded passage, viral infection was detected using real-time RT-PCR, and viral titers were calculated as fluorescent focus units (FFU) per ml as described^38^. In parallel, the cells and supernatants were prepared for transmission electron microscopy as described elsewhere^12^.

### Serologic analysis

Recombinant nucleoprotein (N protein) of SGLV was used in ELISA antibody assay. N gene of SGLV (GenBank accession number MT328777) was cloned into pET-30a (Novagen), and the resulting recombinant plasmid was transformed into *Escherichia coli* BL21 DE3 strain. Bacteria were grown at 37°C for 3 h before induction with 1 mM IPTG for 16 h. The recombinant N protein was purified using Ni^2+^ resin (Thermo Fisher Scientific, USA). An indirect ELISA assay was conducted with the recombinant N protein as the coating antigen (3 ng/μl) to evaluate viral specific antibodies in patient serum. The negative control contains eight pools of serum samples from 100 healthy individuals. The cut off value of the reaction was calculated as the mean OD of negative control sera plus three standard deviations.

For the microneutralization test, serially diluted serum samples were mixed with an equal volume of 100 FFU virus (100 μl), respectively. After incubation at 37°C for 1.5 h, the mixture was added to a 96-well plate with SMMC-7721 cells in quadruplicate. The plate was incubated in 5% CO_2_ at 37°C for 4 days. Viral infection was detected by using indirect immunofluorescence assay (IFA). The end-point titer was expressed as the reciprocal of the highest dilution of serum that prevented infection.

### Statistics analysis

All statistical analyses were performed with GraphPad Prism software version 5.0 (GraphPad Software Inc., San Diego, USA). The differences between each sample group were compared using the student *t* test. The prevalence of virus infection in pooled ticks were calculated by maximum likelihood estimation (MLE) using the program PooledInfRate^39^. The differences were considered statistically significant at *p*-value less than 0.05.

### Ethics statement

The study was reviewed and approved by the ethics committee of the Inner Mongolia General Forestry Hospital (2016001) in accordance with the medical research regulations of China. All participants provided written informed consent.

## Author contributions

Conceptualization was provided by Q.L., J.Q. and W.W. The methodology was developed by J.M., Z.-D.W., L.L., J.-W.S., and L.-N.L. Investigations were carried out by J.M., X.Z., Z.-D.W., L.L., J.-W.S., C.C., Y.-H.Z., L.S. and L.-N.L. The original draft of the manuscript was written by J.M., X.Z. and Z.-D.W. Review and editing of the manuscript were carried out by Q.L., J.Q., and W.W. Funding acquisition was performed by Q.L. and J.Q. Resources were provided by X.-L.L., S.-Z.H., H.-T.S., L.-X.M. and Z.-L.C. Q.L. provided supervision.

## Acknowledgments

We thank Prof Jing-Fei Wang and Dr Shi-Da Wang at Harbin Veterinary Research Institute, Chinese Academy of Agricultural Sciences for electron microscopic analysis of virus. This work was supported by the National Key R&D Program of China (2017YFD0501700) and the National Natural Science Foundation of China (31672542 and 31372430), the Pearl River Talent Recruitment Program in Guangdong Province of China (2019CX01N111), and the “Three Major” Scientific Research Projects of Sun Yat-Sen University in 2020 (50000-31143406).

